# Insights into the activation of Kinesin1 from the molecular characterisation of JIP3/4 binding to Kif5b

**DOI:** 10.1101/2022.09.09.507386

**Authors:** Fernando Vilela, Mélanie Chenon, Christophe Velours, Jessica Andreani, Paola Llinas, Julie Ménétrey

**Affiliations:** Université Paris-Saclay, CEA, CNRS, Institute for Integrative Biology of the Cell (I2BC), 91198, Gif-sur-Yvette, France; Institut Pasteur, Université Paris Cité, CNRS UMR3528, Structural Image Analysis Unit, F-75015 Paris, France; CBM-UPR4301, Rue Charles Sadron, 45071 ORLEANS Cedex 2; Fundamental Microbiology and Pathogenicity Laboratory, UMR 5234 CNRS-University of Bordeaux, SFR TransBioMed, 33076 Bordeaux, France

**Keywords:** JIP3, JIP4, RH1 domain, Kif5, Kinesin1

## Abstract

Whereas our understanding of kinesin auto-inhibition mechanisms is improving faster, important insights into kinesin activation mechanisms such as those controlled by cargo-motor adaptors are still missing. JIP3 and JIP4 are versatile motor-cargo adaptors for kinesin1 and dynein-dynactin motors enabling bi-directional transport on microtubules. JIP3 activates kinesin1 heavy chains, independently of kinesin1 light chains. In this report, we characterize the molecular details of the binding of the kinesin1 heavy chain, Kif5b to the motor-cargo adaptors, JIP3 and JIP4, using biophysical approaches. The definition of the exact binding site of Kif5b, as well as the specificity of interaction between JIP3 and JIP4 provide new insights into kinesin1 activation.

## INTRODUCTION

Kinesin1 is a molecular motor that transport cargos, such as vesicles, organelles, mRNA and protein assemblies along microtubules (MTs) throughout the cell. When not transporting cargo, kinesin1 is maintained in an autoinhibited state, which prevents futile ATP consumption and congestion on MTs. Activation of kinesin1 is controlled by the binding of both MT-associated proteins and motor-cargo adaptors (Cross and Dodding, 2019; Hooikaas et al., 2019). Recruitment of a single (Watt et al., 2015) or a coordination of two motor-cargo adaptors (Blasius et al., 2007) can trigger kinesin1 activation. Although we know a lot about kinesin1 inhibition, we still lack important insights concerning the regulating mechanisms such as the structural basis of this motor activation by cargo-motor adaptors.

JIP3 and JIP4 (JNK-Interacting Protein 3 and 4) are close paralogs identified as MT-based motor-cargo adaptors (Bowman et al., 2000; Cavalli et al., 2005). JIP3 is mainly found in neurons and has emerged as a regulator of axonal transport (Edwards et al., 2013; Rafiq et al., 2022; Sun et al., 2013; Watt et al., 2015), while JIP4 is ubiquitously expressed and involved in organelle positioning (Boecker et al., 2021; Willett et al., 2017), endosomal membrane reshaping (Marchesin et al., 2015) and recycling (Montagnac et al., 2011). Such functions in intracellular transport are based on the ability of JIP3 and JIP4 (JIP3/4) to coordinate kinesin1 (Bowman et al., 2000; Nguyen et al., 2005; Sun et al., 2011) and dynactin/dynein motors (Cavalli et al., 2005; Montagnac et al., 2009) motors enabling bi-directional transport on MTs. Furthermore, JIP3 activates kinesin1, enhancing its motility along MTs (Sun et al., 2011).

Kinesin1 is an heterotetramer composed of a homodimer of heavy chains (KHC, with the three paralogs Kif5a-c in human) associated to two light chains (KLC; four paralogs, KLC1-4). KHC consists of the N-terminal motor domain (MD) followed by a long stalk formed by several coiled-coils and the C-terminal Tail encompassing a coiled-coil region and an unstructured region named, the distal tail. Kinesin1 interacts with JIP3/4 through a dual mode of binding: the Tail region of KHC binds to the N-terminal region of JIP3/4 (Sun et al., 2011), while KLC binds to the middle part of JIP3/4 (Nguyen et al., 2005). Direct interaction of JIP3 with KHC, independently of KLC, is functional and enhances the motility of kinesin1 along MTs (Sun et al., 2011; Watt et al., 2015). The motility of KHC is regulated *via* direct intramolecular interactions between the MD and the distal tail. In short, KHC folds back enabling the interaction of a short motif within the distal tail, called the Isoleucine-Alanine-Lysine (IAK) motif, with the MD. This interaction prevents structural rearrangements required for MT-binding and the ATPase cycle (Cai and Verhey, 2007; Dietrich et al., 2008; Hackney and Stock, 2000; Kaan et al., 2011; Seiler et al., 2000). KLC is also involved in the autoinhibition of kinesin1 (Vale and Friedman, 1999; Verhey et al., 1998), a recent study reported that KLC inhibits kinesin1-MT interaction independently from the proposed intramolecular interaction within KHC (Chiba et al., 2022). Whereas the binding of JIP3 to KLC2 has been characterized structurally (Cockburn et al., 2018), such molecular details are not available for the KHC:JIP3/4 interaction. These data are required to understand how this interaction enables kinesin1 activation

Here, we present the molecular characterization of JIP3 and JIP4 binding to Kif5b using purified proteins. Our biophysical experiments define the minimal binding site of Kif5b and JIP3/4 for the interaction and provide critical information on the specificity of Kif5b for JIP3 and JIP4. Altogether, our data provide new insights into KHC activation by JIP3.

## RESULTS AND DISCUSSION

### Biophysical characterization of Kif5b-Tail and JIP3/4-Nter fragments

In order to perform biophysical studies on JIP3/4 binding to Kif5b and define the minimal binding sites of both proteins, we designed various fragments of the Tail region of Kif5b (Kif5b-Tail) and of the N-terminal part of JIP3 and JIP4 (JIP3/4-Nter), and validated their structural integrity using Circular Dichroism (CD), Analytical UltraCentrifugation (AUC) and/or Multi-Angle Light Scattering (MALS), The Tail region of Kif5b consists of the last part of the cc2 coiled-coil (namely cc2b) encompassing the KLC-binding site, followed by the cc3 coiled-coil and the 50-aa unstructured distal tail (dt). We conceived three fragments of the Tail region of human Kif5b (Fig.1A and Table S1): (i) the cc2b coiled-coil followed by the cc3 coiled-coil and the distal tail (cc2b-cc3-dt fragment), (ii) a truncated form lacking the distal tail (cc2b-cc3 fragment) and (iii) the cc3 coiled-coil alone (cc3 fragment). CD experiments showed that the cc2b-cc3 and cc3 fragments are mainly alpha-helical, while for the cc2b-cc3-dt fragment, the turn/random content increases significantly, consistent with the presence of the unstructured distal tail (see details in Supplementary information). Furthermore, AUC experiments indicate that the three fragments are mainly dimers in solution (data not shown). The N-terminal part of JIP3/4 encompasses an RH1 (RILP-Homology 1) domain which assembles as a dimer of two anti-parallel helices forming a four-helix bundle. The RH1 domain is flanked by a short unstructured region in its N-terminus (N) and followed by a long Leucine Zipper coiled-coil region (LZI) (Fig.1C). For both JIP3 and JIP4, we used a previously characterized fragment that encompasses the RH1 domain followed by the entire LZI coiled-coil (RH1-LZI fragment) (Vilela et al., 2019), as well as three new fragments (Figure 1C and Table S2). The first one contains the N-terminus region (N-RH1-LZI fragment) and the two others were truncated after the third heptad repeat of LZI coiled-coil (RH1-lz3 and N-RH1-lz3 fragments). MALS experiments demonstrated that the presence of the N-terminus, as well as the truncation of the last eleven heptads of the LZI have no impact on the dimeric form of JIP3 and JIP4 (see details in Supplementary information). Previously, we showed that the RH1 domain is not stable in the absence of the LZI coiled-coil (Vilela et al., 2019). Our new results indicate that the first three heptad repeats of LZI are sufficient to stabilise the dimeric form of JIP3 and JIP4. Collectively, these results showed that all Kif5b-Tail fragments and JIP3/4-Nter fragments used here, exhibit the expected structural integrity required for binding assays.

**Figure 1.**
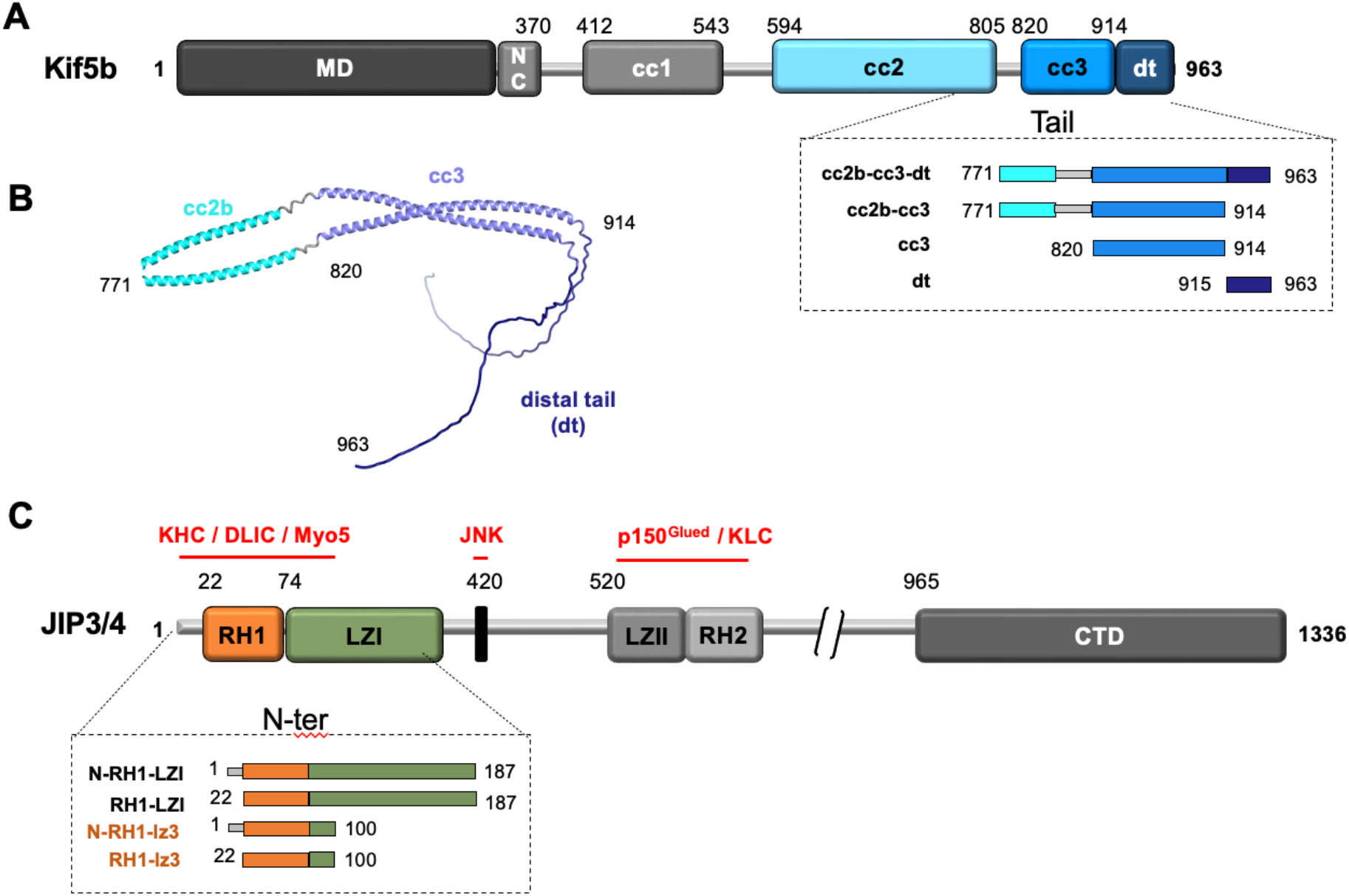
Structural integrity of Kif5b-Tail and JIP3/4-Nter fragments. **(A)** Schema of the full length Kif5b. MD: Motor Domain; NC: Neck Coil; cc: coiled-coil; dt: distal tail. Sequence limits of human Kif5b are indicated at the beginning of the known domains. The dash lined-box (bottom) shows the fragments of the C-terminal part of Kif5b used in this study. **(B)** Model of Kif5b-cc2b-cc3-dt using the AlphaFold2-based pipeline ColabFold with the AlphaFold-Multimer version (Evans et al., 2022; Jumper et al., 2021; Mirdita et al., 2022). **(C)** Schema of the full length JIP3/4 proteins. The JIP3/4 partners are indicated in red. The dash lined-box (bottom) shows the fragments of the N-terminal part of JIP3/4 used in this study.

### The cc3 coiled-coil of Kif5b is the binding site for JIP3

To characterize the mode of binding of JIP3 to Kif5b, we used the different fragments of both proteins to analyse their interaction in solution by Microscale Thermophoresis (MST). First, we confirmed that both proteins interact together using the longest Kif5b-Tail and JIP3-Nter fragments. The Kif5b-cc2b-cc3-dt fragment binds to the JIP3-N-RH1-LZI fragment with a Kd of 18.2 ± 2.9 nM (Fig.2A, Table 1). When the N-terminus of JIP3 is removed (RH1-LZI fragment), the Kd is 3.7-fold higher than that of the N-RH1-LZI fragment (Fig.2A and Table 1) indicating a role, although minor, for this unstructured region in the interaction. However, whether the N-terminus is directly in contact with Kif5b or is involved in the stabilisation of the RH1 domain of JIP3/4 remains to be determined. When the LZI is truncated after the third heptad repeat (N-RH1-lz3 fragment), the Kd is the same as that of the N-RH1-LZI fragment (Fig.2A and Table 1) revealing no role of the full LZI in the interaction. Therefore, our data identify the RH1-lz3 region of JIP3 as the main binding site for Kif5b. This is consistent with previous data defining a minimal Kif5-binding site to residues 50 to 80, which overlaps to the RH1 domain (Sun et al., 2011).

**Figure 2.**
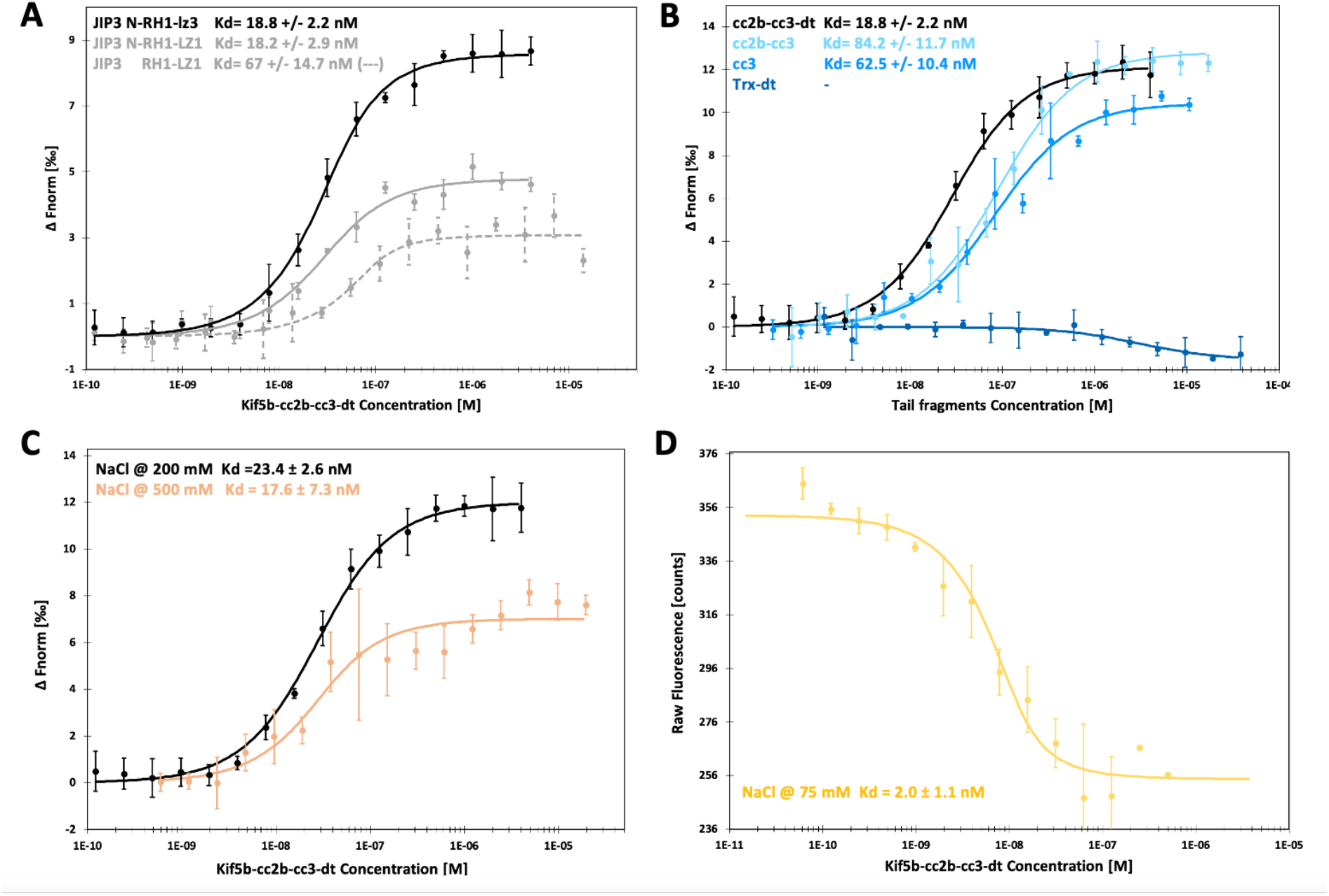
MST characterization of the interaction between Kif5b-Tail and JIP3-Nter. **(A)** Superposition of the dose-response curves from the titration of JIP3-N-RH1-LZ1 (grey line), JIP3-RH1-LZ1 (grey dashes) and JIP3-N-RH1-lz3 (black) with KIF5b-cc2-cc3-dt. The concentration of the labelled JIP3-Nter fragments was kept constant at 20 nM, while the concentration of the KIF5b-cc2-cc3-dt was varied between to 1.2.10^-4^ to 4 uM. **(B)** Superposition of the dose-response curves from the titration of labelled JIP3-N-RH1-lz3 with Kif5b-Tail fragments: cc2b-cc3-dt (black), cc2b-cc3 (cyan), cc3 (light blue) and Trx-dt (dark blue). The concentration of the labelled JIP3-N-RH1-lz3 was kept constant at 20 nM while the concentration of the tail fragments was varied from 5.2.10^-4^ to 17 uM (kif5b-cc2b-cc3), 3.2.10^-4^ to 10.6 uM (Kif5b-cc3) and 1.2.10^-4^ to 4 uM (Kif5b-cc2b-cc3-dt). **(C)** Superposition of the dose-response curves from the titration of JIP3-N-RH1-lz3 with KIF5b-cc2-cc3-dt at different ionic strength: 200 mM NaCl (black) and 500 mM NaCl (light orange). **(D)** Initial fluorescence dose-response curve from the titration of JIP3-N-RH1-lz3 with KIF5b-cc2-cc3-dt at 75 mM NaCl (yellow). At 75 mM NaCl, there is a fluorescence signal variation due to the interaction so the data must be analyzed using this signal and not the thermophoresis. JIP3-N-RH1-lz3 is negatively charged and KIF5b-cc2-cc3-dt is positively charged, the diminution of NaCl concentration may allow additional interactions in a region near to the fluorophore, that should be masked at higher ionic strength. Error bars indicate the standard deviation of three replicates.

**Table 1.** MST interaction experiments between various KIF5b-Tail fragments and JIP3/4-N-ter fragments.

| Kif5b-Tail | JIP3/4-Nter | NaCl (mM) | Kd (nM) |
| --- | --- | --- | --- |
| Screening JIP3 fragments |  |  |  |
| cc2b-cc3-dt | N-RH1-LZI | 200 | 18.2 ± 2.9 |
|  | RH1-LZ1 |  | 67.0 ± 14.7 |
|  | N-RH1-lz3 |  | 18.8 ± 2.2* |
| Screening Kif5b fragments |  |  |  |
| cc2b-cc3-dt | N-RH1-lz3 | 200 | 18.8 ± 2.2* |
| cc2b-cc3 |  |  | 84.2 ± 11.7 |
| cc3 |  |  | 62.5 ± 19.4 |
| dt |  |  | - |
| Screening of NaCl Concentration |  |  |  |
| cc2b-cc3-dt | N-RH1-lz3 | 75 | 2.0 ± 1.1# |
|  |  | 200 | 18.8 ± 2.2* |
|  |  | 500 | 33.0 ± 14.4 |
| Screening JIP4 fragments |  |  |  |
| cc2b-cc3-dt | N-RH1-lz3 | 200 | 758.6 ± 159.2 |
|  | RH1-lz3 |  | 704.7 ± 192.1 |
| Screening JIP3-N-RH1-lz3 mutants |  |  |  |
| cc2b-cc3-dt | H45G-C46R | 200 | 167.0 ± 29.8 |
|  | N61A |  | 20.0 ± 5.7 |
|  | L70F-S71A-E72Q-N73D |  | 50.2 ± 4.5 |
|  | E77Q |  | 42.0± 7.8 |

Buffer: 50 mM Tris pH 8, 10 mM MgCl<sub>2</sub>, 0.05% tween 20 and NaCl (concentration indicated in the table). \* Same measurement. #Kd measured from initial fluorescence dose-response curves.

On the other hand, to define the minimal region of Kif5b required for JIP3 binding, the interaction between the N-RH1-lz3 fragment of JIP3 and truncated Kif5b-Tail fragments were analysed and compared. When the distal tail is removed, Kif5b shows a 4.5-fold decrease in binding affinity (cc2b-cc3-dt vs cc2b-cc3; Fig.2B and Table 1). However, no significant interaction was detected with the distal tail alone at the tested concentrations (in absence of the cc2b-cc3 part; Fig.2B and Table 1). These data indicate that the distal tail might play a minor role in JIP3 binding, possibly in stabilising the cc3 coiled-coil or the JIP3-Kif5b assembly. When the cc2b part is removed, Kif5b exhibits similar binding affinity for JIP3 (cc2b-cc3 vs cc3; Fig.2b and Table 1), showing that the cc2b region is not involved in JIP3 binding. Altogether, this interaction analysis defines the cc3 region of Kif5b as the main binding site for JIP3. Thus, these results indicate that the RH1-lz3 domain of JIP3/4 makes direct interaction with the cc3 region of Kif5b and that two unstructured regions from JIP3 (N-terminus) and Kif5b (distal tail) have minor impacts on this interaction, possibly in stabilizing the proteins alone or their complex.

Finally, the physicochemical forces responsible for the interaction were assessed by analysing the binding between the JIP3-N-RH1-lz3 and the Kif5b-cc2b-cc3-dt fragments at three different concentrations of NaCl. No significant difference is observed for the binding affinity between 200 mM and 500mM NaCl, but a 10-fold difference is observed when saline concentration is reduced to 75 mM NaCl (Fig.2C and 2D and Table 1). These results show that the interaction between JIP3-Nter and cc3 has a major hydrophobic component, as it is maintained at 500 mM NaCl. The increase in the affinity at lower ionic strength indicates the formation of electrostatic interactions also, consistent with the electrostatic properties of both JIP3-N-RH1-lz3 (pI 4.8) and the Kif5b-cc2b-cc3-dt (pI 10.2) fragments.

To summarize, this study showed that JIP3 bind to the cc3 coiled-coil of Kif5b. The cc3 region of Kif5 is a well-known binding site for cargos or adaptors, especially those using coiled-coil regions for the interaction, like the SNARE proteins SNAP25 and SNAP23, the ribosome receptor p180, the Glutamate-receptor-interacting protein GRIP1, the Milton/TRAK adaptor and the tropomyosin (Diefenbach et al., 2002; Diefenbach et al., 2004; Dimitrova-Paternoga et al., 2021; Glater et al., 2006; Setou et al., 2002).

### Kif5b exhibits higher affinity for JIP3 than JIP4

Because JIP3 and JIP4 have some sequence differences in the Kif5-binding site (Fig.3A), we wondered if Kif5b may exhibit binding specificity between the two paralogs. MST binding assays showed that Kif5b-cc2b-cc3-dt fragment binds to JIP4-N-RH1-lz3 fragment with a Kd of 760 ± 160 nM, which indicates a 40-fold lower affinity compared to the equivalent JIP3 fragment (Fig. 3B and Table S2). This result points to a higher specificity of Kif5b for JIP3 compared to JIP4. However, it should be noted that the Kd of the JIP4:KHC interaction is similar to what was measured for motor light chains (KLC (Cockburn et al., 2018) or DLIC (Celestino et al., 2022)) bound to JIP3/4. Consistently, functional studies showed that JIP4 is recruited by kinesin1 heavy chain and is involved in kinesin-based transport process (Sato et al., 2015; Suzuki et al., 2020). Thus, we suggest that such a binding affinity difference between JIP3 and JIP4 could account in variations of kif5b regulation mechanisms, such as motor activation.

**Figure 3.**
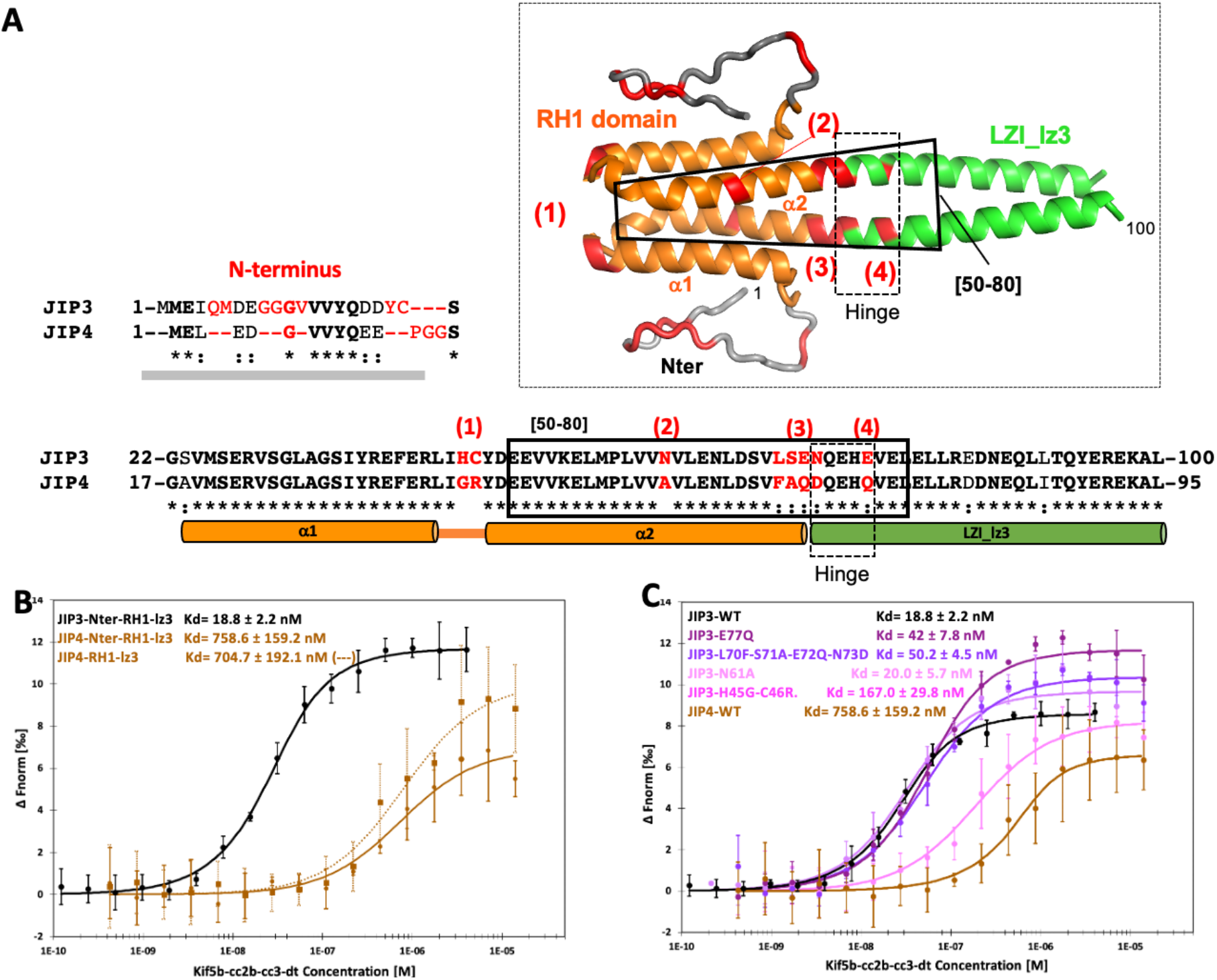
Differences in the interaction of the N-terminal region of JIP3 and JIP4 with the tail of Kif5b analyzed by microscale thermophoresis (MST). **(A)** Sequence comparison of the N-terminal region of JIP3 and JIP4 with a schematic representation of the folding of this region. (Inset) Model of JIP3-N-RH1-lz3 fragment using the AlphaFold2-based pipeline ColabFold with the AlphaFold-Multimer version (Evans et al., 2022; Jumper et al., 2021; Mirdita et al., 2022). Sequence differences between JIP3 and JIP4 are shown in “red” and correspond to the mutations studied in this work (B and C). The region of JIP3-Nter corresponding to residues 50-80 is indicated by a bold black square and the hinge region between the RH1 domain and the LZI region by a dash-line black square. **(B)** Superposition of the dose-response curves from the titration of labelled JIP3-N-RH1-lz3 (black), JIP4-N-RH1-lz3 (brown dots) and JIP4-RH1-lz3 (brown) with Kif5b-cc2-cc3-dt. The concentration of the labelled JIP3 and JIP4 fragments was kept constant at 20 nM while the concentration of KIF5b-cc2-cc3-dt was varied between 1.2.10^-4^ to 16 uM. **(C)** Superposition of the dose-response curves from the titration of labelled JIP3-N-RH1-lz3 and the JIP3 to JIP4 mutants with Kif5b-cc2-cc3-dt.

To analyse further the structural basis of this specificity, we investigated the impact of sequence differences between JIP3 and JIP4 on the binding affinity for Kif5b using truncated fragments and site-directed mutants. Sequence differences are observed in the N-terminus, in the RH1 domain especially into the loop that connect the α1 and α2 helices (α1-α2 loop), the α2 helix, as well as in the hinge region between the RH1 domain and the LZI and at the beginning of the LZI (Fig. 3A.). First, the N-terminus region diverges with several small insertions in JIP3 compared to JIP4 (Fig. 3A). The JIP4-RH1-lz3 fragment which lacks the N-terminus binds to the Kif5b-cc2b-cc3-dt fragment with a Kd of 705 ± 192 nM which is equivalent to that measured for the JIP4-N-RH1-lz3 fragment (Fig.3B and Table 1), indicating that the N-terminus of JIP4 has no impact on Kif5b binding. This result diverges from that of JIP3 for which the truncation of the N-terminus leads to a 3.7-fold binding difference (Fig.2A and Table 1). Thus, the sequence differences between JIP3 and JIP4 at the N-terminus contribute, albeit in a minor way, to the binding specificity of Kif5b for JIP3/4. Then, site-directed mutagenesis was performed on the JIP3-N-RH1-lz3 fragment in order to change for JIP4 sequence (Fig.3A): (1) His45-Cys46 were mutated to Gly-Arg (H45G-C46R), (2) Asn61 to Ala (N61A), (3) Leu-Ser-Glu-Asn (70-73) to Phe-Ala-Gln-Asp (L70F-S71A-E72Q-N73D) and (4) Glu77 to Gln (E77Q). MST experiments were performed to determine the binding affinity of these four JIP3-N-RH1-lz3 mutants for Kif5b-cc2b-cc3-dt (Fig.3C and Table 1). Data showed that there is no difference in affinity between the wild-type JIP3-N-RH1-lz3 fragment and the N61A mutant. A recent report showed that the V60Q mutation of JIP3 does not prevent the interaction with the mouse Kif5c (Celestino et al., 2022). This confirms that the region 60-61 of JIP3 is not involved in Kif5 binding. Then, only a 2- to 3-fold difference in affinity is observed for the E77Q and L70F-S71A-E72Q-N73D mutants, and a 9-fold difference for the H45G-C46R mutant. However, whether these residues are in direct contact with Kif5b or are involved in the stabilisation of JIP3/4 structure remains to be determined. Thus, our data show that sequence differences in the α1-α2 loop are the main contributions for the higher affinity of Kif5b for JIP3, while those in the N-terminus and the α2 helix contribute only marginally.

### Does JIP3 binding to Kif5b induce structural rearrangements?

In order to detail the structural basis of JIP3 recruitment by Kif5b, we tried to predict a model of the Kif5b:JIP3 complex using AlphaFold2 (Evans et al., 2022; Jumper et al., 2021; Mirdita et al., 2022). However, despite varying the boundaries of the Kif5b and JIP3 sequences, the stoichiometry of the complex, the multiple sequence alignments used for prediction and the AlphaFold parameters (monomer or multimer version), we did not converge on a reliable model. The difficulties to predict a structural model for Kif5b:JIP3 complex could be explained by the coiled-coil regions involved in the interaction. Coiled-coils are versatile structural motifs (Truebestein and Leonard, 2016) that can change their multimeric arrangement upon partner binding, as shown for instance by the recent structural study of the actin binding protein tropomyosin 1 (*a*Tm1) bound to the cc3 region of Khc (Dimitrova-Paternoga et al., 2021). The Khc-binding region of *a*Tm1 consists of a dimeric coiled-coil that dissociates in presence of Khc-cc3 and a single helix associates with the Khc-cc3 dimeric coiled-coil to form a tripartite coiled-coil complex (Dimitrova-Paternoga et al., 2021). Another type of structural rearrangement could be a change in the register of the coiled-coils, as was observed in the dynein stalk domain and the dynein activator BicD2 (Carter et al., 2008; Liu et al., 2013). Further work will be necessary to decipher the structural basis of the binding mode of Kif5b to JIP3/4.

### Insights into the activation of KHC by JIP3

The characterization of JIP3 binding to Kif5b presented in this study provides new insights into KHC activation by JIP3, independently of KLC binding. Our data have shown that JIP3 is recruited by the cc3 part of KHC, with no direct interaction with the distal tail. Therefore, the auto-inhibitory IAK motif of Kif5b, embedded in the distal tail, does not contribute to the interaction with JIP3. Thus, our results exclude a model in which JIP3 would release the MD:Tail interaction by competing with the MD for the IAK motif binding, and rather favour an allosteric model. One possibility would be a proximity effect between the JIP3:KHC-cc3 and the MD:IAK assemblies triggering steric hindrance. Another, non-exclusive, allosteric model for KHC activation might be due to structural rearrangements in the cc3 part of KHC upon JIP3 binding that will affect the distal tail region, leading to IAK motif dissociation from the MD. Further work will be required to characterize the allosteric mechanism of KHC activation by JIP3, especially by studying differences between JIP3 and JIP4 on this mechanism.

## MATERIALS AND METHODS

### Microscale Thermophoresis

JIP3-Nter fragments were labelled using the Protein Labelling Kit Blue-NHS (NanoTemper Technologies). The labelling reaction was performed according to the manufacturer’s instructions in the supplied labelling buffer applying a concentration of 20 μM protein (molar dye:protein ratio ≈ 3:1) at room temperature for 30 min. The labelled protein fragments were stored in MST buffer (20 mM Hepes pH 7, 150 mM NaCl). For the binding assays, the labelled JIP3-Nter fragments were adjusted to 20 nM with an assay buffer containing 50 mM Tris pH 8.0, 200 mM NaCl, 5 mM MgCl2, 0.05 % Tween 20. Kif5b fragments were buffer exchanged to the assay buffer and a series of 16 1:1 dilution was prepared. For the measurement, each Kif5b fragment dilution was mixed with one volume of labelled JIP3-Nter fragments. The samples were loaded into Monolith NT.115 Premium Capillaries MO-K025 (NanoTemper Technologies). MST was measured using a Monolith NT. 115 instrument (NanoTemper Technologies) at 25°C. Instrument parameters were adjusted to 40 % LED power and low MST power. Data of three independent replicates were analysed using the signal from an MST-on time at 1.5-2.5 s (MO.Affinity Analysis software version 2.3, NanoTemper Technologies).

## Acknowledgments

We thank Drs. Philippe Chavrier, Kristen Verhey and Reza Khayat for the gift of the JIP3, JIP4 and Kif5b plasmids, respectively. This work has benefited from the expertise of Magali Aumont-Nicaise and the Macromolecular interaction measurements Platform of I2BC supported by the French Infrastructure for Integrated Structural Biology (FRISBI) ANR-10-INSB-05. JA is supported by the Agence Nationale de la Recherche through grant ESPRINet (ANR-18-CE45-0005-01). We are grateful to Drs. Davy Martin and Human Rezaei (INRA, VIM-UR0892, Jouy-en-Josas) for access to their CD equipment. Advice given by Pierre Soule (NanoTemper Technologies GmbH) have been a great help for MST experiments. We thank Dr. Benoît Gigant for useful discussions, as well as for careful reading and comments on the manuscript.

